# Loss of hemagglutination ability by H3N2 influenza A virus, subclade K

**DOI:** 10.64898/2026.07.20.739523

**Authors:** Ruonan Liang, Pascal Lexmond, Oliver C. Grant, Mark Pronk, Roland J. Pieters, Ron A.M. Fouchier, Geert-Jan Boons, Björn F. Koel, Robert P. de Vries

## Abstract

Seasonal human H3N2 influenza viruses, subclade K (J.2.4.1), have been the predominant influenza A viruses in the Northern hemisphere influenza season of 2025/2026. Since 2024, the vaccine virus A/Darwin/6/21 has emerged in different antigenic variants. Antigenic changes are frequently caused by amino acid substitutions near the hemagglutinin (HA) receptor-binding pocket, which can also affect receptor binding properties, such as hemagglutination. Hemagglutination is crucial for assessing antigenicity using the hemagglutination inhibition (HAI) assay, and a loss of binding to turkey erythrocytes could significantly hamper this process. In this study, we explored how substitutions in or around the HA receptor-binding site affect binding to glycans at the molecular level. We employed ELISA, glycan array, flow cytometry, hemagglutination assays, and tissue staining. Substitutions at positions 140, 192, and 223 establish clade J viruses that emerged in 2024. Computational analysis of HA in complex with an elongated glycan reveals that mutation F192 forms a CH-Pi interaction to stabilize the binding. Based on this background, substitutions in antigenic sites A and B within subclade K viruses exhibit a binding preference for elongated glycans, which are not displayed on turkey erythrocytes. Conversely, our previously established glyco-remodeled erythrocytes are efficiently bound by these subclade K H3N2 viruses and could support influenza surveillance and vaccine development.

## Introduction

Influenza A virus (IAV) continues to pose a significant burden on public health, causing seasonal epidemics and pandemics. IAV contains two surface glycoproteins, hemagglutinin (HA) and neuraminidase (NA)^1^. HA binds to cell-surface glycans terminating in sialic acid (SIA), mediates membrane fusion during viral entry, and is the major target of the immune system, while NA cleaves the SIA from viral receptors to release virions.

H3N2 is one of two influenza A virus subtypes circulating in humans. When it was introduced in humans during the 1968 pandemic, the HA receptor specificity switched from α2-3-linked to α2-6-linked sialylated glycans^2,3^. Human respiratory tract tissue^4^ and tracheal ciliated cells^5^ have Asn-linked (N-glycans) with extended branches, terminating with α2-6-linked SIA, a unique characteristic of the airway glycome. They are extended from the mannose core (Man3GlcNAc2Asn) by LacNAc (Galβ1-4GlcNAcβ) sequences and can contain multiple LacNAc repeats. In recent decades, H3N2 viruses have altered their receptor-binding properties through antigenic mutations in response to immune selection pressure ^6,7^, while maintaining functional receptor binding. During this process, H3N2 viruses exhibited different binding specificities between short and long LacNAc N-glycans^8–10^. Recent human H3N2 viruses have shown a preference for long, branched sialoglycans with tri-LacNAc repeats^11,12^. However, after 2021, H3N2 strain 3C.2a1b.2a.2 viruses regained the ability to bind SIA presented on a shorter di-LacNAc unit ^13,14^.

The hemagglutination inhibition assay is the cornerstone for monitoring influenza virus antigenicity using turkey erythrocytes^15–17^. During the evolution of H3N2 viruses, many have lost their ability to agglutinate turkey erythrocytes^18,19^. This is because turkey erythrocytes do not display tri-LacNAc-containing structures. Consequently, the loss of erythrocyte binding has made it difficult to select vaccine antigens, and thus guinea pig erythrocytes are now often employed, while they exhibit low HA titers^20^. Glycan-modified erythrocytes, rather than normal erythrocytes, have been shown to overcome this limitation^11^. Additionally, glyco-engineered cells that display extended glycan receptors were created to analyze HA receptor-binding specificity or to enhance the growth of recent H3N2 viruses ^21,22^.

Since February 2025, clade 3C.2a1b.2a.2a.3a.1 is the main H3N2 virus circulating in the human population, and these viruses have been genetically divided into subclades J.1-J.4^23^ and their subsequent HA descendants, J.2.3 (N158K and K189R), J.2.4 (T135K and K189R), and J.2.5 (S145N and K189R), circulating in different continuities. Since the winter of 2025, J.2.4.1 (renamed subclade K), characterized by J.2.4 subclade, carrying additional mutations K2N, S144N, N158D, I160K, Q173R, T328A, and S378N, has been the globally dominant H3N2 virus^24^. Post-infection ferret antisera raised against NH2025–2026 vaccine viruses (A/Croatia/10136RV/2023 and A/District of Columbia/27/2023) showed poor recognition of emerging A(H3N2) subclade K viruses, whereas antisera raised against the updated subclade K vaccine strain A/Darwin/1415/2025 showed improved recognition; consequently, WHO updated the recommended vaccine composition for the NH2026–2027 season^24^.

Receptor binding avidity of HA correlates with antigenic drift^25^. To understand such phenotypic changes of subclade K at the molecular level, we introduced mutations near or within receptor-binding sites of recent H3N2 virus HA proteins. We examined the receptor binding of circulating H3N2 viruses using ELISA, FACS, tissue staining, glycan arrays, and glyco-engineered turkey erythrocytes. Subclade K clearly prefers binding to elongated N-glycans; thus, it is unable to hemagglutinate turkey erythrocytes. Our study supports the use of glycoengineering of turkey erythrocytes for antigenic characterization of influenza viruses, including the currently circulating subclade K viruses.

## Results

### I223V reduced, while I192F rescued binding to Di-LacNAc N-glycans

Representative vaccine strains recommended by the WHO for the 2022-2027 northern hemisphere winter influenza seasons were selected for sequence alignment in this study (Fig. S1). Our sequence alignments showed that since clade 3C.2a1b.2a2a.3a.1 emerged, the mutations I140K, I192F, and I223V have appeared in the receptor binding site.

To assess how these single mutations affect glycan-binding specificity and avidity, a quantitative enzyme-linked immunosorbent assay (ELISA) was performed using a set of biotinylated glycans^14^. They were symmetrical biantennary N-glycans with mono-, di-, and tri-LacNAc repeats and asymmetrical compounds with di- and tri-LacNAc repeats on the α3 arm. Amino acid substitutions I140K, I192F, and I223V were introduced in A/Darwin/6/21 (Fig. S1). A/Darwin/6/21 wildtype (WT) HA bound efficiently to compounds **C** symmetrical α2–6 sialylated di-LacNAc glycan, **D** asymmetrical sialylated tri-LacNAc at the Manα3 arm, and **E** symmetrical sialylated tri-LacNAc biantennary N-glycan(Fig. 1A), which is consistent with our previous study. Compared with WT, I140K and I192F exhibited lower EC50 values, indicating enhanced binding avidity to compounds **C**, **D,** and **E**. Notably, I192F exhibited the lowest EC50 values for compounds **D** and **E**, suggesting the strongest binding avidity toward these two compounds. In contrast, I223V resulted in a higher EC50 for compounds **C**, **D**, and **E**, indicating reduced avidity for them. Given their positions relative to the receptor binding site, we surmise that I140K and I223V affect affinity for the terminal SIA portion of the glycan receptor, whereas the I192F mutation may interact with the repeating LacNAc portion.

**Fig. 1.**
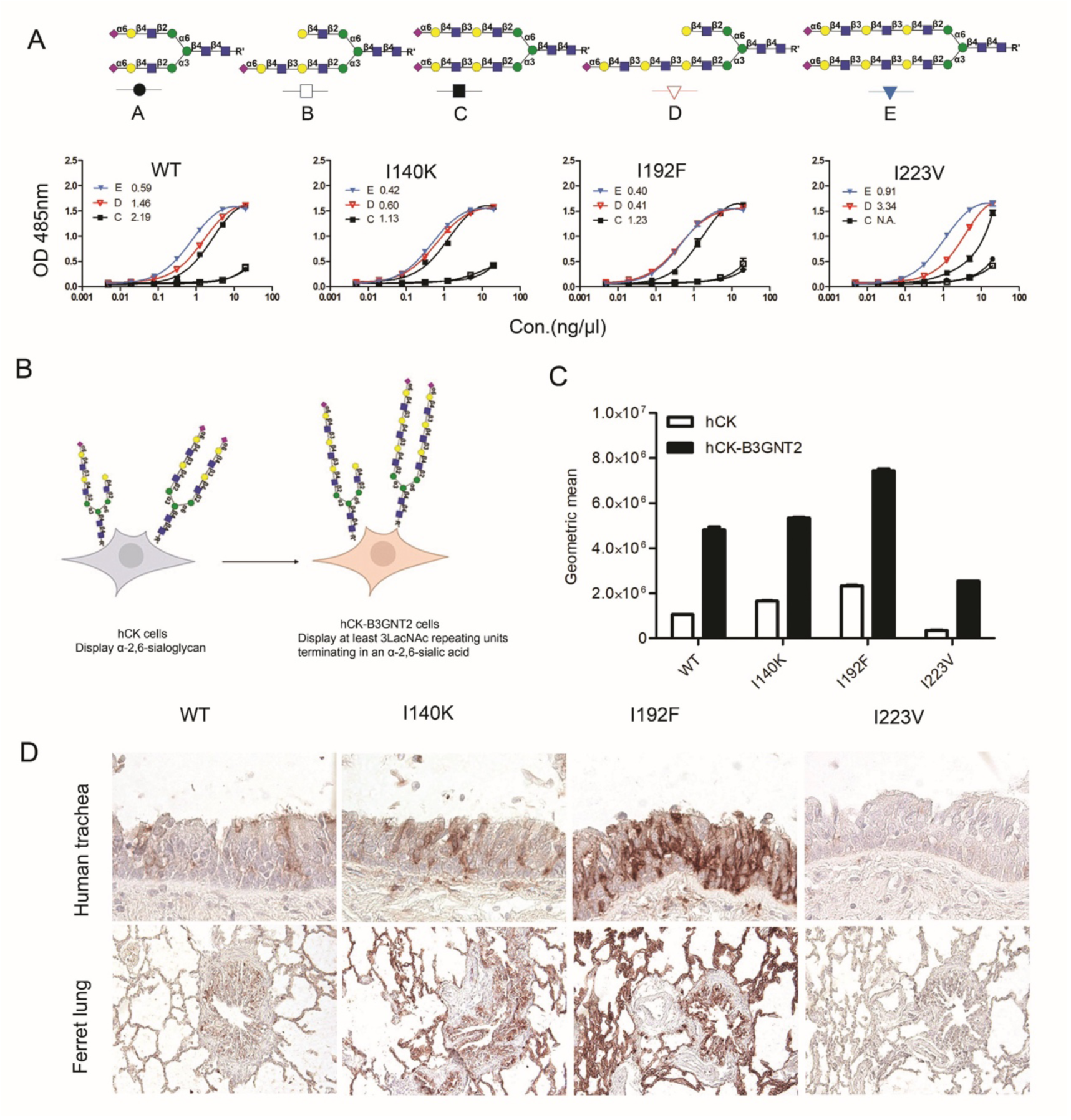
I223V reduced, while I192F rescued binding to Di-LacNAc N-glycans. A. Binding avidity of A/Darwin/6/21 HAs (wt and mutations) to glycans were measured by ELISA (Value represents EC50, meaning the half-maximal effective concentration). B. hCK and hCK-B3GNT2 cells displayed different types of α 2,6-sialylgycans. C. Binding of A/Darwin/6/21 HAs (WT and single mutations) proteins in hCK and hCK-B3GNT2 cells were analyzed by flow cytometry. Binding levels were quantified based on the geometric mean fluorescence intensity. And values on the Y-axis are presented in scientific notation. D.Binding of A/Darwin/6/21 HAs (WT and single mutations) to human trachea and ferret lung were examined by tissue staining. For ELISA and flow cytometry assays, data were obtained from three independent experiments with N=2 or N=3 replicates. Tissue staining experiments were performed in two independent experiments.

To assess the effects of amino acid substitutions on cellular binding in vitro, hCK and hCK-B3GNT2 cells were employed (Fig. 1B)^21^. The engineered hCK-B3GNT2 cells display an increased number of LacNAc repeating units terminated with α 2,6-SIAs compared to hCK cells^26,27^. The binding ability of A/Darwin/6/21 WT and mutated counterparts was tested using flow cytometry (Fig. 1C). A/Darwin/6/21 WT demonstrated a strong preference for hCK-B3GNT2 cells compared to hCK cells. Substitutions I140K had a slightly increasing binding to hCK and hCK B3GNT2, and I192F increased binding to both cell types significantly, especially for the hCK-B3GNT2 cells. Conversely, I223V showed lower binding to both cells, but still preferred the hCK-B3GNT2 cells.

Ferret lung and human tracheal tissues are established models for studying the receptor-binding properties of hemagglutinins. A/Darwin/6/2021 WT bound to the epithelial layer in the human trachea, bronchi, and alveoli in the ferret lung (Fig. 1D). I140K and I192F displayed intense binding signals to these tissues, especially I192F, whereas I223V showed a very weak binding signal to human trachea and ferret lung sections. Thus, using a suite of receptor-binding assays, we demonstrate that I223V reduces binding, while I140K and I192F increase binding.

### A combination of I140K/I192F/I223V form a new background, assuming clade J

A/Massachusetts/18/2022 and A/Thailand/8/2022 were recommended as cell- and egg-based A(H3N2) virus vaccine strains for the 2024-2025 Northern Hemisphere influenza season. Compared with A/Darwin/6/2021, these vaccine strains contain these three HA amino acid substitutions in receptor-binding sites: I140K, I192F, and I223V (Fig. S1). To analyze how these mutations affect binding specificity, different combinations of them were tested (I140K/I192F, I140K/I223V, I192F/I223V, and I140K/I192F/I223V). ELISA analyses showed that, compared with I140K (Fig. 1A), the I140K/I223V exhibited higher EC50 values for compounds **C**, **D**, and **E**, accompanied by a marked rightward shift in the binding curves, indicating reduced binding avidity (Fig. 2A). In contrast, compared with I192F, I192F/I223V retained a preference binding for the elongated glycan **D** and **E**, but exhibited reduced binding avidity to compound **C**, as indicated by its higher EC50 value. This confirmed that the I223V substitution reduces binding avidity (Fig. 1A). I140K/I192F retains a similar binding avidity to the previous single-mutation I140K or I192F. I140K/I192F/I223V (3Muts) exhibited a preference for binding to elongated glycan **D** and **E** compared with glycan **C**. Complementary flow cytometry demonstrated a similar binding trend in binding to hCK-B3GNT2 cells (Fig. 2B), and tissue binding analyses revealed a strong signal in human trachea and ferret lung sections (Fig. 2C).

**Fig. 2.**
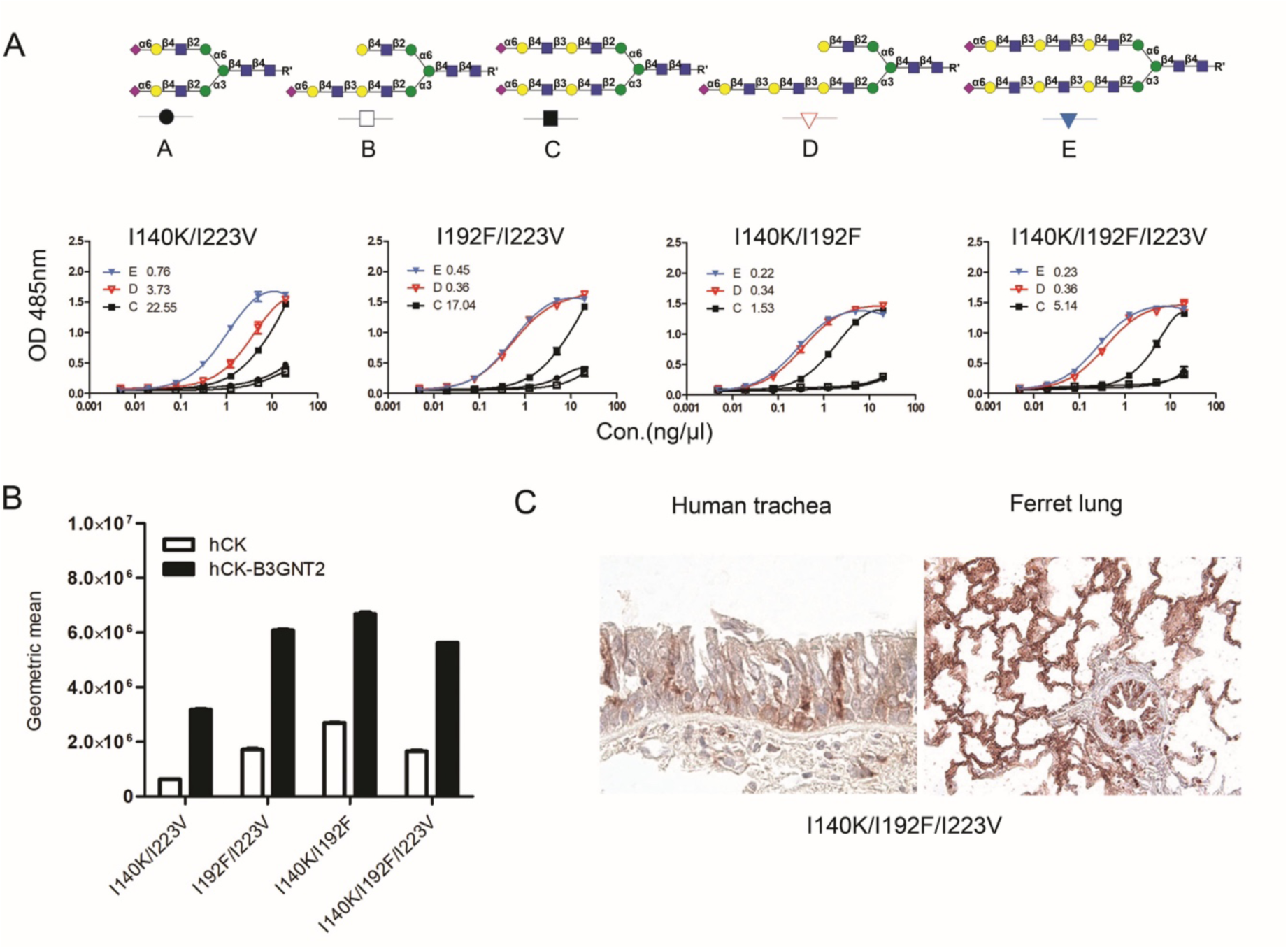
The Combination of I140K/I192F/ I223V formed a background of clade J. A. Binding avidity of A/Darwin/6/21 HAs (wt and different combined mutations) to glycans was measured by ELISA.(Value represents EC50, meaning the half-maximal effective concentration) B. Binding of A/Darwin/6/21 HAs (WT and combined mutations) to hCK and hCK-B3GNT2 cells was analyzed by flow cytometry. C. Binding of A/Darwin/6/21 HAs (WT and mutations) to human trachea and ferret lung was tested by tissue staining. For ELISA and flow cytometry assays, data were obtained from three independent experiments with N=2 or N=3 replicates. Tissue staining experiments were performed in two independent experiments.

### Subclade K showed a preference for elongated N-glycans

Relative to the HA of the A/Massachusetts/18/2022 2024-2025 clade J vaccine strain, subclade K viruses carry amino acids K2N, N122D, T135K, S144N, N158D, I160K, Q173R, K189R, and K276E in HA1(Fig. 3A). Positions 135, 144, 158, 160, and 189 are not only part of the receptor-binding sites but also located in antigenic site A (135 and 144) and site B (158, 160, and 189). Some of them are likely to drive antigenic change during influenza virus evolution^6^, substitutions at positions 158 and 189 have been implicated in an antigenic cluster transition^6^. Subclade K viruses have drifted antigenically and showed reduced cross-reactivity with ferret antisera against 2024-2025 northern hemisphere vaccine strain A/Thailand/08/2022 and A/Massachusetts/18/2022 (J) and 2025-2026 vaccine strain A/Croatia/10136V/2023 (J.2) vaccine^28,29^. However, population-based immunity did reveal efficient protection^30,31^.

**Fig. 3.**
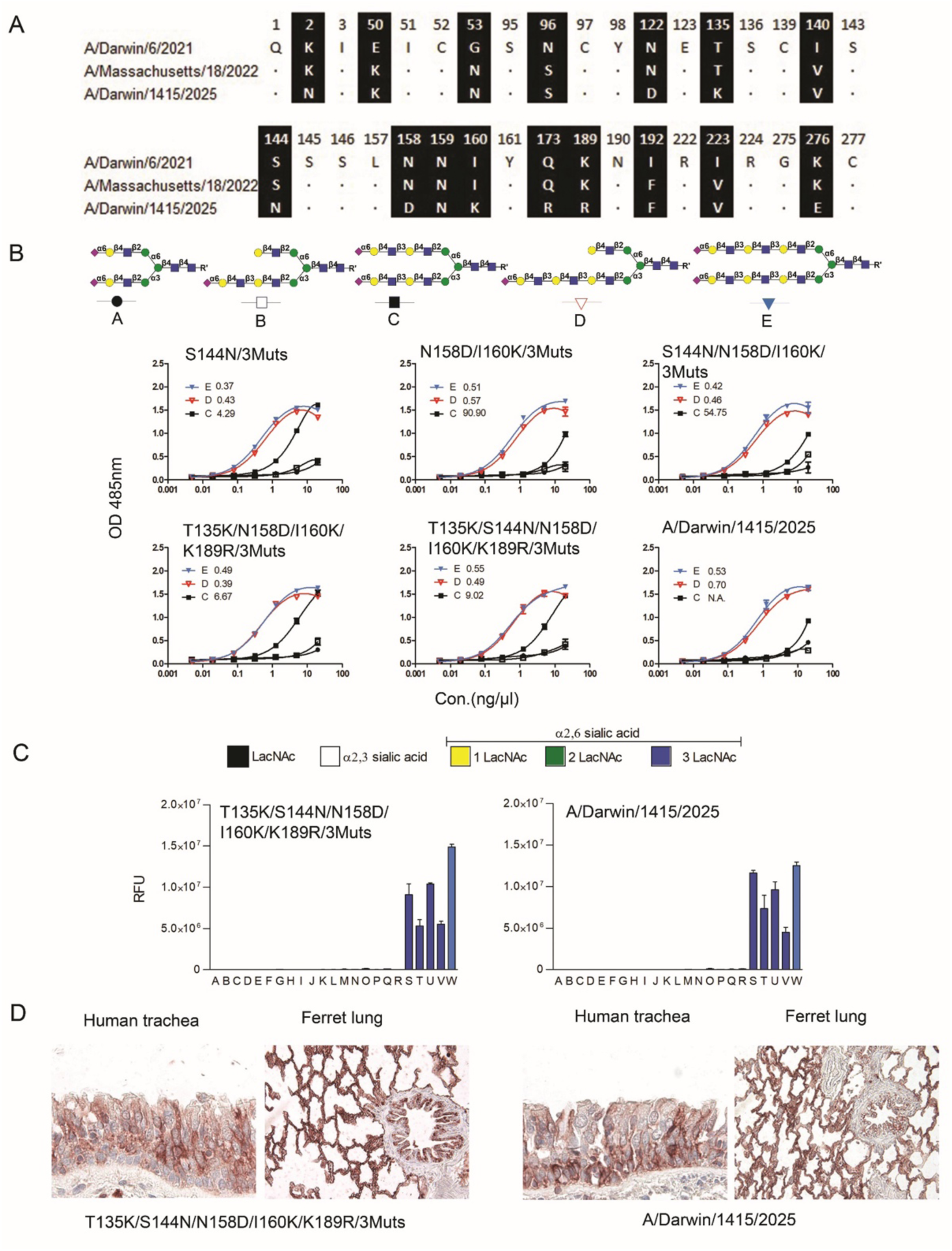
Subclade K showed a preference binding to elongated N-glycans. A. Alignment of A/Darwin/6/2021, A/Massachusetts/18/2022, and A/Darwin/1415/2025. B. Binding avidity of Subclade K and related mutations to glycans was measured by ELISA. All mutations were introduced based on the 3Muts background (Value represents EC50, meaning the half-maximal effective concentration). C.Binding specificity of Subclade K proteins(S144N/T135K/K189R/N158D/I160K/3muts and A/Darwin/1415/2025) was analyzed by glycan array. RFU represents relative fluorescence units. The glycan profile used for the array is shown in Fig. 5A. D. Binding characteristics of subclade K proteins to human trachea and ferret lung were examined by tissue staining. For ELISA assays, data were obtained from three independent experiments with N=2 replicates. Glycan array data were calculated over four replicates. Tissue staining experiments were performed in two independent experiments.

To study the effect of the emerged mutations on receptor-binding properties we introduced them on top of the 3Muts (I140K/I192F/I223V) HA, creating S144N/3Muts, N158D/I160K/3Muts, T135K/K189R/3Muts (J.2.4), S144N/N158D/I160K/3Muts, T135K/ N158D/I160K/ K189R/3Muts, T135K/S144N/ K189R/3Muts, and S144N/T135K/K189R/N158D/I160K/3Muts. However, T135K/K189R/3Muts and S144N/T135K/K189R/3Muts express poorly in HEK293S GnTI(-) cells (Fig. S2). Among the successfully expressed recombinant proteins, S144N/3Muts displayed a similar binding trend to 3Muts (Fig. 3B). In contrast, N158D/I160K/3Muts preferred binding to elongated compounds **D** and **E**, and almost lost binding ability to compound **C**, as indicated by the increased EC50 value. When combining S144N with N158D/I160K, the resulting HA retained a similar binding pattern to that of N158D/I160K/3Muts, suggesting that N158D/I160K predominantly determines the altered glycan-binding preference, whereas the additional S144N substitution has a limited effect on receptor-binding avidity. However, T135K/N158D/I160K/K189R/3Muts displayed an ability to bind compound **C**. The T135K/S144N/N158D/I160K/K189R/3Muts protein contains all substitutions in the RBS of Subclade K (A/Darwin/1415/2025). However when we used the full A/Darwin/1415/2025 HA we observed a significant loss of binding to compound **C**. Indiciating that substitutions outside of the RBS can fine tune receptor binding properties.

To assess binding specificity to a wider range of N-glycans, we employed a glycan array, in which T135K/S144N/N158D/I160K/K189R/3Muts and A/Darwin/1415/2025 exhibited binding to α2,6-linked SIA presented on tri-LacNac repeat-containing N-glycans (Fig. 3C). We previously demonstrated that such specificity correlates with an inability to recognize turkey erythrocytes^11,13^. Both subclade K proteins bound to human tracheal and ferret lung tissues, indicating that structures such as compound **D** are present in these respiratory tissues (Fig. 3D).

### Molecular modeling of A/Darwin/1415/2025 binding to compound B and D

To assess how the different amino acids change receptor binding properties we generated 3D structures of compound **B** and **D** in complex with A/Darwin/1415/2025 and A/Darwin/6/2021 and performed molecular dynamics simulations. In multiple replicate molecular dynamics simulations of A/Darwin/1415/2025, the aromatic ring of the I192F mutation spontaneously formed a stable, long-lasting CH-pi stacking interaction with the 3rd Galactose residue from the SIA terminal that is uniquely present in tri-LacNAc-containing glycans such as glycan **D** (Fig. 4 B and D and Tables S1). In the shorter, di-LacNAc glycan **B** the 192F instead interacts with the tri-mannose of the N-glycan core (Fig. 4 A and C). A per-residue MMGBSA calculation indicates that this interaction is weaker, which correlates with the experimental observation that I192F dramatically enhances affinity for extended poly-LacNAc glycans, and re-establishes A/Darwin/6/2021 level binding to tri-LacNAc containing glycans as observed in our suite of receptor binding assays.

**Fig. 4.**
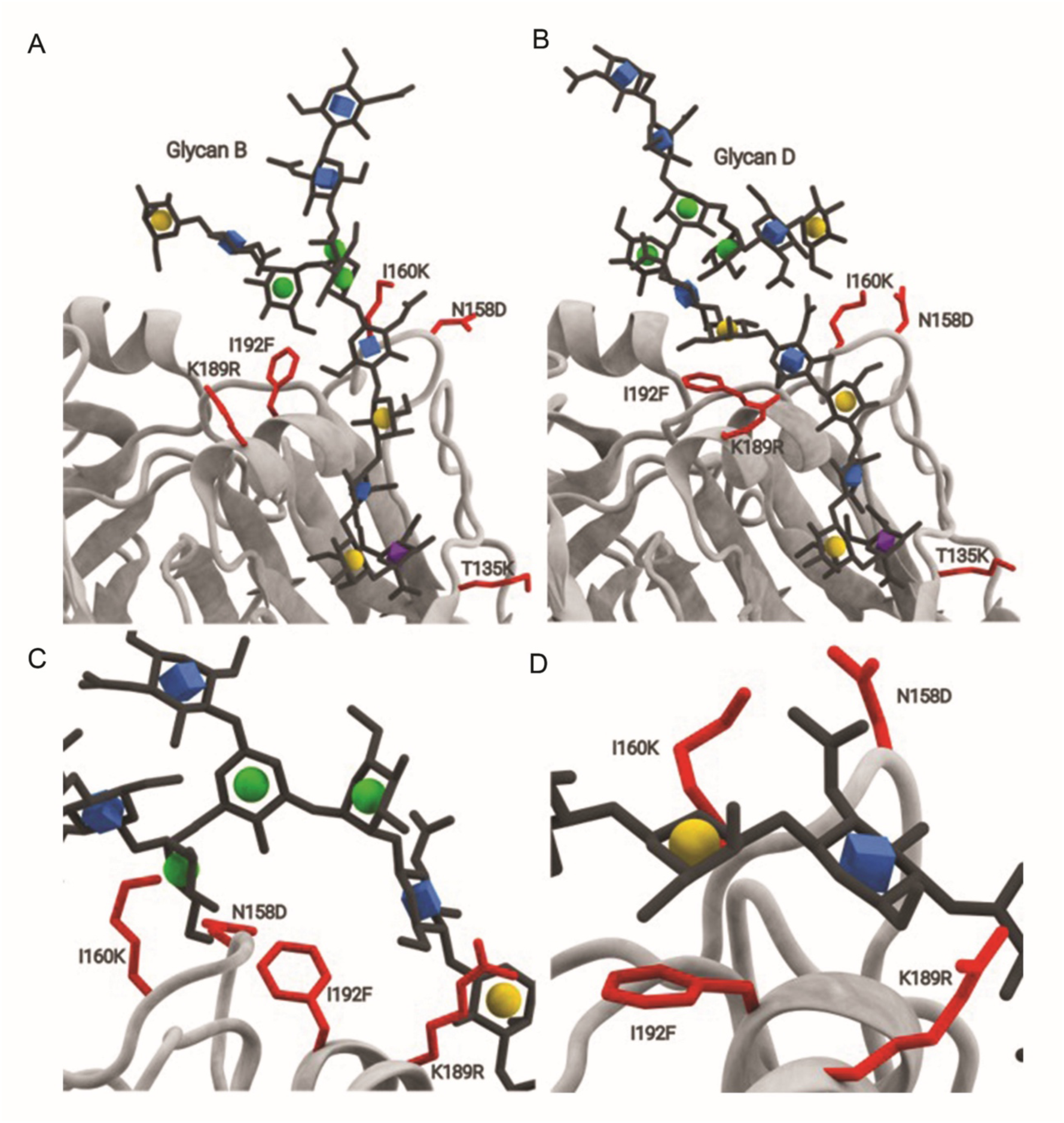
Molecular modeling of Darwin/1415/2025 (grey ribbon) demonstrates unique binding to array glycans B and D (shown as stick with SNFG symbols in the ring). A. The tri-mannose core of the shorter Glycan B interacts with I192F. B. The 3rd Galactose residue, uniquely present in tri-lacNAc containing glycans such as glycan D, forms a very stable stacking interaction with I192F that spontaneously forms in each of the five replicate simulations and is stably present throughout the simulation.

### Receptor binding specificities of subclade K viruses

Our recombinant protein approach demonstrates that subclade K has strict specificity for elong ated N-glycans. To confirm this observation, we elected to employ full H3N2 viruses and examined A/NL/02082/2025 as an example of clade J 2.2, which additionally carries substitutions S124N and R222K in the receptor binding site compared with clade J, and A/NL/02209/2025 and A/NL/02145/2025, representing subclade K. A/NL/02082/2025 displayed receptor specificity to a wide range of N-glycans displaying α2,6-linked SIA presented on di-LacNac repeats as indicated by the green bars (Fig. 5A). This virus was also able to bind a mono-LacNAc containing N-glycan (**#M**). Conversely, the two viruses representing subclade K, reverted to a strict specificity to α2,6-linked SIA presented on tri-LacNAc repeat-containing N-glycans. These same viruses were analyzed for hemagglutination properties, in which the subclade J.2.2 virus readily hemagglutinates turkey erythrocytes, whereas the subclade K viruses fail to do so (Fig. 5B). We have previously presented a platform for creating glyco-engineered erythrocytes to facilitate antigenic characterization of H3N2 viruses that fail to bind turkey erythrocytes^11,32^. We treated turkey erythrocytes with a sialidase and resialylated them with hST6G1 (α2,6 Sia-LN). Another batch was treated with B3GNT2 and B4GALT1 during desialylation and thereafter treated with hST6G1, creating glyco-engineered erythrocytes α2,6 SIA poly-LN. α2,6 SIA poly-LN erythrocytes were agglutinated by a pre-2020 genetic clade 3C.2a1 A/NL/1797/2017 virus. Whereas the A/NL/384/2019 (genetic clade 3C.3a) agglutinated untreated, α2.6-Sia, and α2,6-Sia-poly-LN turkey erythrocytes(Fig. 5B), as previously demonstrated^11^. A/NL/02082/2025 hemagglutinates all types of erythrocytes, including turkey and α2,6-Sia-LN versions, as expected given its broad receptor binding specificity in the glycan array. On the other hand, both A/NL/02209/2025 and A/NL/02145/2025 subclade K viruses hemagglutinated α2,6-Sia-poly-LN to identical titers. This data illustrates that subclade K viruses revert to binding only poly-LacNAc-containing N-glycans. Additionally, guinea pig erythrocytes were employed to assess the agglutination properties of subclade K, which displayed a hemagglutination titer of only 8 (Fig. 5B). This value was approximately 3- to 4-fold lower than that obtained using remodeled turkey erythrocytes.

**Fig. 5.**
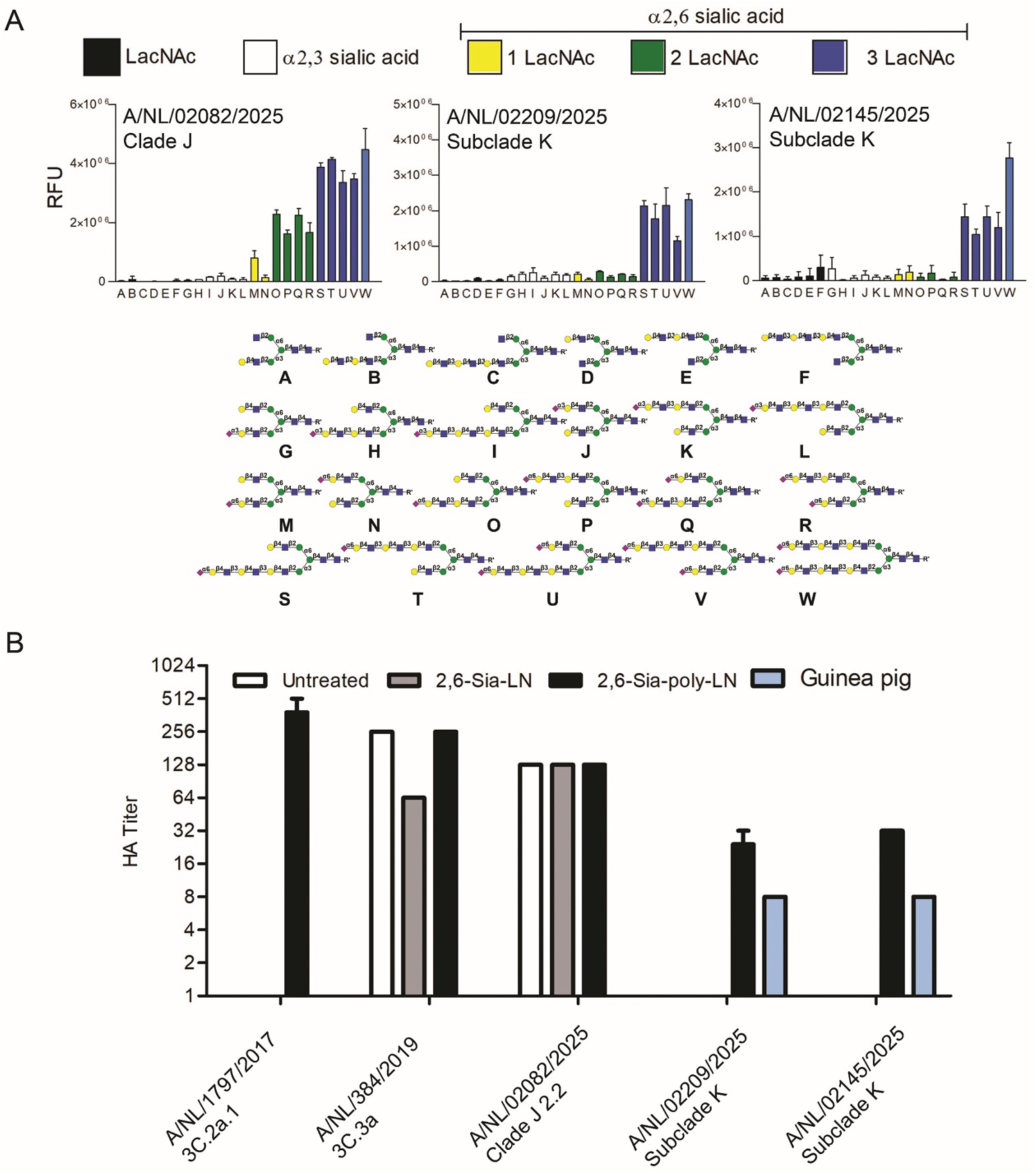
Receptor binding specificities of subclade K. A. Binding specificity of Viruses A/NL/02082/2025, A/NL02209/2025, and A/NL/02145/2025 was analyzed by glycan microarrays. And beneath is the glycan profile imprinted on microarray slides. B. Hemagglutination ability was performed by HA assay using pig erythrocytes. HA assay test of these untreated, α2,6 Sia poly-LN and α2,6 Sia poly-LN remodified turkey erythrocytes and guinea erythrocytes. Glycan array data were calculated over four replicates. HA assay data were analyzed in duplicate.

## Discussion

Here, we demonstrate that specific substitutions in antigenic sites A and B of subclade K HA result in the inability to agglutinate turkey erythrocytes. As part of the HA receptor-binding site, the 220-loop is the most conserved region and may interact with amino acids in the 190-helix. The related epistasis between the 190-helix and 220-loop has been implicated in the adaptation of human H1 and H3 subtypes^33,34^. And they are primarily involved in substitutions at positions 190-225. Although substitutions at position 223 have not previously been reported to affect receptor binding properties, our finding suggests that this residue may contribute to altered receptor interactions. The I192F substitution on the surface of the RBS, which affects receptor binding, has not been observed in H3N2 viruses but has been identified in H9N2 viruses^35^.

The N-linked glycosylation site (Asn-X-Ser/Thr) at position 158 has been shown to affect receptor binding properties ^36,37^. A new glycosylation site at position 144^38^ appeared not to affect receptor binding in our hands, suggesting that not all glycosylation sites have equivalent effects on receptor binding. Although there is no glycosylation site at position 158 in contemporary H3N2 viruses, other substitutions in this 158-160 sequon affect receptor binding by directly interacting with LacNAc^14,39,40^. Similarly, I192F contributes to a receptor-binding mode through direct interaction, illustrating how glycan flexibility is employed for binding.

The T135K/S144N/K158D/I160K/K189R/3Muts HA protein showed a binding avidity to symmetrical α2,6-sialilated di-LacNAc compound **C**, whereas the full A/Darwin/1415/2025 did not. The difference here are mutations outside the RBS, such as N96S, N122D, and Q173R, which thus clearly further fine-tune receptor binding. Previous research also showed that a compensatory mutation in H1 changed the HA binding ability to SIA receptors on the cell surface^41^.

As turkey erythrocytes are poorly bound by many human H3N2 viruses in the last two decades, guinea pig erythrocytes have been adopted as an alternative, as they display glycans containing extended LacNAc repeat structures with mono- or di-sialylation^10^. However, the HA titers obtained with guinea pig erythrocytes remain generally low^11^, and these erythrocytes are not preferred because they are smaller due to their mammalian origin and are more difficult to interpret in routine assays. Nevertheless, some studies have successfully employed guinea pig erythrocytes in HAI assays to antigenically characterize subclade K, using prior vaccines during antigenic drift^30,31^. However, viruses with low binding avidity are easily inhibited in the HAI assa^25^, which may explain why seasonal vaccines still exhibit substantial HAI activity against subclade K.

Given the poor agglutination of certain erythrocyte types, HA assays may fail to provide reliable virus detection or an accurate virus titer^42^. Notably, when an HA titer is very low, the guinea pig erythrocytes do not compensate for intrinsically weak HA-receptor binding, limiting the reliability of HA-based readouts for both virus titration and antigenic characterization^42^. Our previous study reported no or very low HA titers for A/NL/1797/2017 and A/NL/384/2019^11^. Together with our current findings, this suggests that guinea pig erythrocytes may not be suitable for antigenic characterization or for assessing vaccine efficacy, as their use may lead to inaccurate evaluation of viral antigenicity.

Other methods of antigenically characterizing influenza A viruses do exist^43,44^, and their development is urgently needed. However, all alternatives so far are more laborious and expensive, and less straightforward and benchmarked than the HAI assay^17^. Glycoengineering of turkey erythrocytes provides a means to measure the antigenicity of subclade K viruses by HAI. Subclade K is not unique in its inability to hemagglutinate turkey erythrocytes, as viruses before 2020^11^ and recent viruses circulating in the Netherlands^32^ also failed to agglutinate these cells. With ongoing viral evolution, glyco-remodeled erythrocytes could be used for antigenic surveillance of both human H1N1 and H3N2 viruses.

## Material and methods

### Protein expression and purification

HA encoding cDNAs of A/Darwin/6/21was cloned into pCD5 expression vectors as described previously^45^. HAs mutations were made using site-directed mutagenesis and checked by Sanger sequencing. These cDNAs were cloned in frame with a secretion signal sequence, the Twin-strep(WSHPQFEKGGGSGGGSWSHPQFEK); IBA, Germany), a GCN4 trimerization domain (RMKQIEDKIEEIESKQKKIENEIARIKK), and morange2. All the HAs were expressed in HEK293S GnTI(-) and purified from cell culture supernatant as described previously. Transfection was performed using DNA and polyethyleneimine I in a ratio 1:8. After 5-6h, the transfection medium was replaced by 293 SFM II expression medium (Gibco), supplemented with sodium bicarbonate (3.7 g/L), Primatone RL-UF (3.0 g/L, Kerry, NY, USA), glucose (2.0 g/L), glutaMAX (1 %, Gibco), valproic acid (0.4 g/L), and DMSO (1.5 %). At 5 days after transfection, cell culture supernatants were collected, and Strep-Tactin sepharose beads (IBA, Germany) were used to purify the HA proteins according to the manufacturer’s instructions.

### Phylogenetic tree

The entire length of the hemagglutinin protein sequences (all the selected H3N2 data are the 2022-2027 northern hemisphere vaccine strains) were downloaded from the GISAID platform^38^. followed by alignment by MAFFT. The phylogenetic tree was constructed using MAFFT.

### Enzyme-linked immunosorbent assay

ELISA was performed as described previously^14^. Briefly, Nunc Maxisorp 96-well plates (Thermo Scientific, #442404) were coated with 50 μL of 5 ug/mL streptavidin in PBS overnight at 4 ℃ followed by blocking with 300 μL of 1% BSA in PBS-T for 3h at room temperature. Then, streptavidin-coated plates were coated with 50 μL of 50 nM compounds in PBS overnight at 4 ℃, followed by blocking with 1% BSA in PBS-T again. HAs at 20 μg/ml were precomplexed with strepmab and goat-anti-human antibodies (Invitrogen, #31410) in a 1:0.65:0.325 molar ratio on ice for 30 min. Precomplexed proteins were added to the plates, diluted serially 1:3, and incubated for 90 min at room temperature. It was developed by 50 ul of OPD buffer and stopped by 25 ul of 2.5 M H_2_SO_4_ after 5 min. The UV reader (Polar Star Omega, BMG Labtech) measured the optical density at 485 nm. The data were calculated from three independent experiments in duplicate. The binding avidity of HA to glycans was quantified by determining the half-maximal effective concentration (EC50), defined as the concentration of HA required to achieve 50% of the maximal binding signal. Lower EC50 values indicate stronger apparent binding avidity between HA and glycans.

### Flow cytometry analysis

Cells (hCK and hCK-B3GNT2) were treated with 2 mL TrypLE express enzyme. Cell pellets were harvested after centrifuge and resuspend in FACS buffer (PBS containing 1% FCS and 2 mM EDTA). They were kept on ice before using. 10 μg/mL hemagglutinin was precomplexed with strepmab and goat-anti-human Alexa Fluor 488 in a 1:0.65:0.325 molar ratio on ice for 30 min. The precomplexed proteins were incubated with 50,000 hCK or hCK-B3GNT2 cells for 30 min at 4°C in the dark. Cells were washed with FACS buffer and followed by the centrifuge at 300 rcf for 5 min. The cells were fixed with 1% Paraformaldehyde for 10 min at 4°C in the dark. After being washed with FACS buffer, the cells were resuspended in 100 μL of FACS buffer. Flow cytometry was performed using the CytoFLEX LX (Beckman Coulter). Data was analyzed using FlowJo software. All cells, single cells, and live cells were gated. Mean fluorescence values for triplicates were averaged, and standard deviations were calculated.

### Histochemical tissue staining

Ferret lung were obtained from the division of pathology, department of biomolecular health sciences, faculty of veterinary medicine of Utrecht University. Human trachea were obtained from UMC Utrecht, Department of Pathology, Utrecht, the Netherlands (TCBio-number 22-599). We embedded all the tissues in paraffin. Histochemical tissue staining was performed as previously described. In brief, tissue sections were deparaffinized and rehydrated. The endogenous peroxidase was inactivated by 1% H2O2 in MEOH for 30 min at room temperature. Tissues were blocked by the 3% BSA in PBS overnight at 4°C. The precomplexed HA, strepmab, and goat-anti-human antibodies were incubated with tissues for 90 min at RT. The concentration of HA was 20 μg/ml for human trachea and 50 μg/ml for ferret lung tissues. After incubation, the tissue was developed by AEC substrate for HRP (Abcam) and subsequently stained by hematoxylin.

### Hemagglutination assay

Guinea pig erythrocytes were obtained from Inotiv (Batch NL260608F). Turkey erythrocytes were supplied by Erasmus Medical Center. Turkey erythrocytes were remodified with different lengths of LacNAc, terminal α2,6 SIA, as described previously^11^. 20 μg/mL Hemagglutinin was incubated with strepmab and goat-anti-human HRP on ice for 30 min (same ratio as ELISA). Pre-complexation was diluted serially in 2-fold.) 1% Erythrocytes were added and incubated at RT for 3–4 hours. It was repeated twice in duplicate.

### Glycan microarray binding study

The glycan array microarray was performed as described previously^11,14^. HAs or viruses were precomplexed with human anti-streptag-HRP and goat anti-human-Alexa 555 antibodies in a 4:2:1 molar ratio in 40 μL of PBS-T on ice for 30 min. Subsequently, they were incubated on the glycan array surface in a humidified chamber for 90 min. Then, the slides were washed in PBS-T, PBS, and deionized water. After the water had been removed by centrifugation, the slides were immediately scanned. The analysis of glycan array data was performed using the carbohydrate-microarray processing script^11^. After removing the highest and lowest values, the data was calculated over four replicates.

### Molecular dynamics simulations

A 3D structure for H3/Darwin/6/2021 was available (PDBID: 9BDF)^46^. This structure was mutated to match the vaccine reference strain of Darwin/6/2021 and Darwin/1415/2025 using UCSF Chimera. Sialic acid ligands were extracted from a A/Shandong/9/1993(H3N2) structure (PDBID 8TJ7)^47^ and aligned into the two Darwin structures using UCSF Chimera’s secondary structure sequence alignment tool. The structures for array glycans B and D were generated using GLYCAM-Web^48^ and superimposed onto the crystal structure ligands via the matching GlcNAc unit, resulting in four systems (glycans B and D in Darwin/6/2021 and Darwin/1415/2025). The glycoprotein builder (v1.5.1) available within gmml2 (https://github.com/GLYCAM-Web/gmml2) and on GLYCAM-Web^48^, was used to add Man5Gn2 glycans to each *N-*glycosylation site. The structures were prepared for energy minimization and MD simulation with the tleap module of AmberTools. An octahedral box of Tip5p waters was placed around the glycoprotein, with an 8 Å-edge buffer and counterions to neutralize the system. The GLYCAM_06j-1 force field was used for the glycans and the AMBERff19SB force field for the protein and ions. Five replicates of the initial structure were then subjected to MD simulations using an established 9-step equilibration protocol with a Langevin thermostat and a 50 ns production run, yielding a cumulative simulation time of 250 ns per system.

### Processing and analysis

*MD trajectory postprocessing:* The mmpbsa module of AmberTools was used to calculate the per-residue interaction energy between the glycan ligand and H3 receptor.

## Acknowledgements

R.L. is supported by a CSC fellowship (202209120001). Research reported in this publication was supported by the National Institute of Allergy and Infectious Diseases of the National Institutes of Health under award no. R01 AI165692 (to G.-J.B.). The content is solely the responsibility of the authors and does not necessarily represent the official views of the National Institutes of Health.

---

**Table S1.** A per-residue MMGBSA calculation performed and averaged across the five replicate MD simulations for each systems.

| **Virus** | **Glycan** | **Replicate number** | **AA** | **DeltaG(Kcal/mol)** | **Std Dev** | **Std Err** |
| --- | --- | --- | --- | --- | --- | --- |
| A/Darwin/6/2021 | B | 1 | ILE 192 | -0,5 | 1,1 | 0,1 |
| A/Darwin/6/2021 | B | 2 | ILE 192 | -0,6 | 0,6 | 0,1 |
| A/Darwin/6/2021 | B | 3 | ILE 192 | -0,5 | 0,6 | 0,1 |
| A/Darwin/6/2021 | B | 4 | ILE 192 | -0,5 | 0,8 | 0,1 |
| A/Darwin/6/2021 | B | 5 | ILE 192 | -0,1 | 0,3 | 0,0 |
| A/Darwin/6/2021 | D | 1 | ILE 192 | -1,3 | 0,6 | 0,1 |
| A/Darwin/6/2021 | D | 2 | ILE 192 | -1,2 | 0,7 | 0,1 |
| A/Darwin/6/2021 | D | 3 | ILE 192 | -0,3 | 0,4 | 0,0 |
| A/Darwin/6/2021 | D | 4 | ILE 192 | -1,3 | 0,8 | 0,1 |
| A/Darwin/6/2021 | D | 5 | ILE 192 | -0,8 | 0,6 | 0,1 |
| A/Darwin/1415/2025 | B | 1 | PHE 192 | -1,6 | 0,6 | 0,1 |
| A/Darwin/1415/2025 | B | 2 | PHE 192 | -1,5 | 0,7 | 0,1 |
| A/Darwin/1415/2025 | B | 3 | PHE 192 | -1,8 | 0,8 | 0,1 |
| A/Darwin/1415/2025 | B | 4 | PHE 192 | -2,2 | 0,7 | 0,1 |
| A/Darwin/1415/2025 | B | 5 | PHE 192 | -1,8 | 0,8 | 0,1 |
| A/Darwin/1415/2025 | D | 1 | PHE 192 | -3,1 | 1,3 | 0,1 |
| A/Darwin/1415/2025 | D | 2 | PHE 192 | -4,0 | 1,0 | 0,1 |
| A/Darwin/1415/2025 | D | 3 | PHE 192 | -4,6 | 0,8 | 0,1 |
| A/Darwin/1415/2025 | D | 4 | PHE 192 | -3,4 | 0,9 | 0,1 |
| A/Darwin/1415/2025 | D | 5 | PHE 192 | -3,3 | 0,7 | 0,1 |

| **Summary** | **Glycan** | **AA** | **DeltaG Average** | **Std Dev** |
| --- | --- | --- | --- | --- |
| A/Darwin/6/2021 | B | ILE 192 | -0,44 | 0,20 |
| A/Darwin/6/2021 | D | ILE 192 | -0,97 | 0,42 |
| A/Darwin/1415/2025 | B | PHE 192 | -1,77 | 0,27 |
| A/Darwin/1415/2025 | D | PHE 192 | -3,67 | 0,61 |

**Fig. S1.**
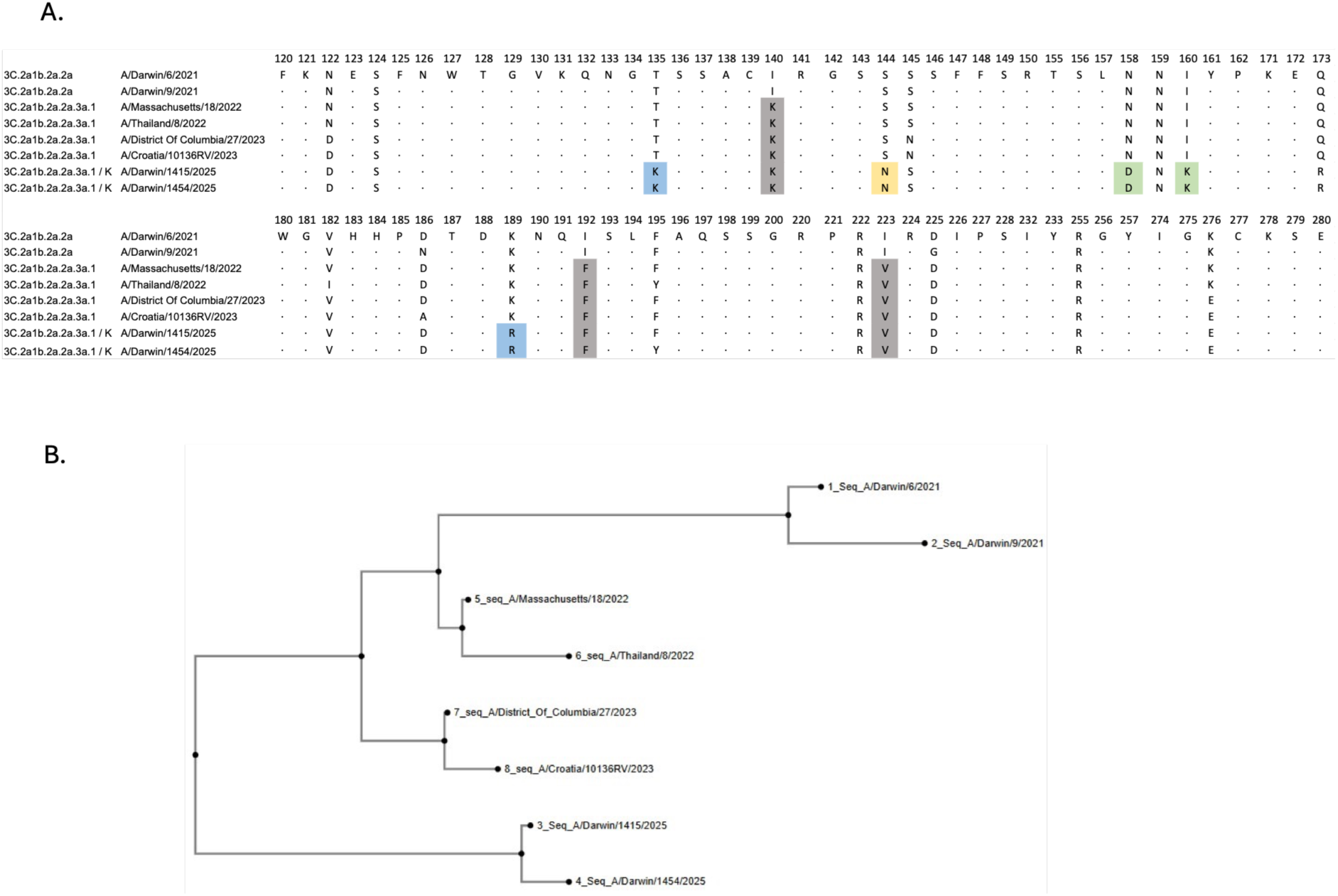
Alignment and Phylogenetic tree in influenza H3N2 strain. A. Sequence Alignment and B. Phylogenetic tree of selected H3N2 virus vaccines during the 2022-2026 northern hemisphere winter influenza seasons

**Fig. S2.**
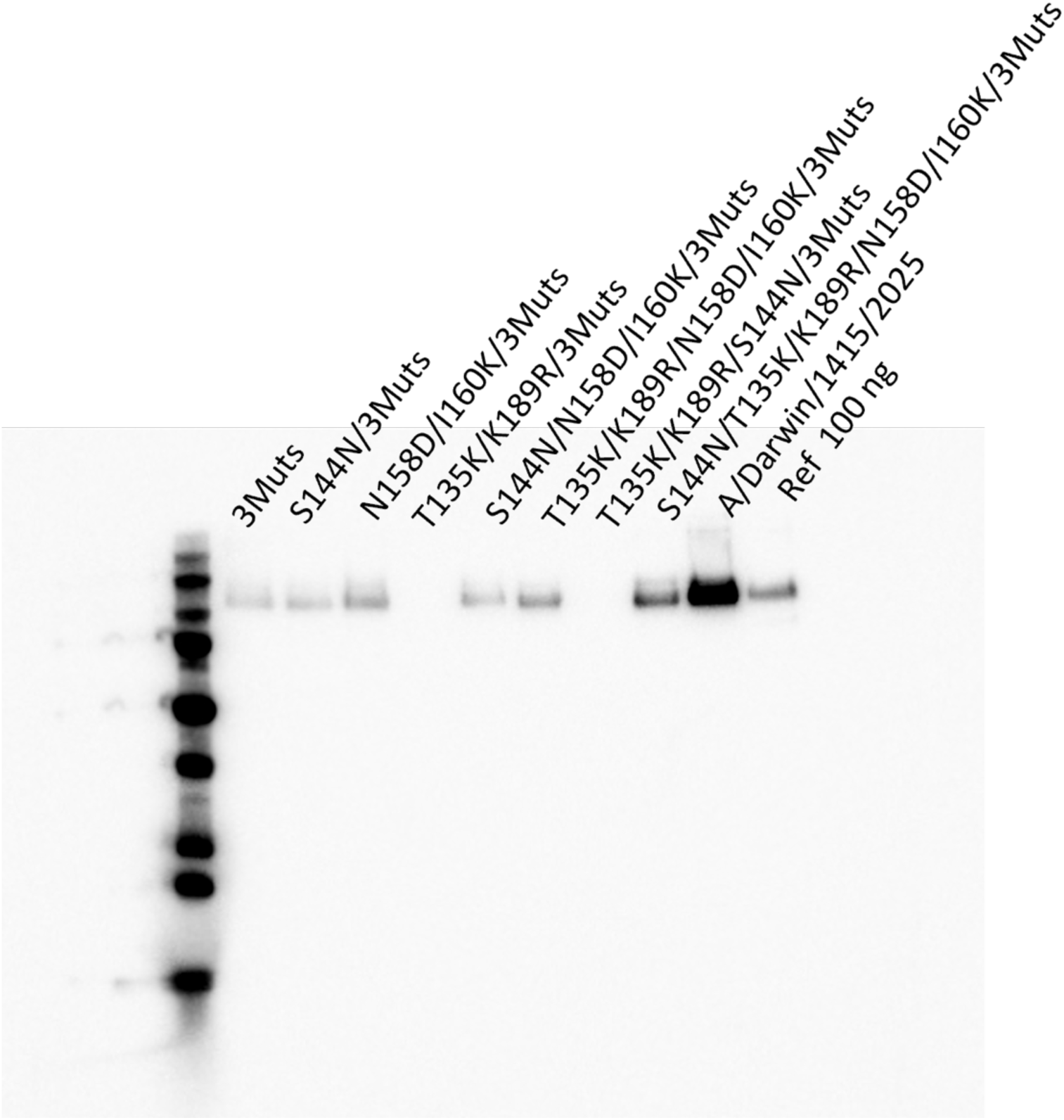
Western blotting of HEK293S GnTI(-) cell supernatants transfected with plasmids containing different mutation combinations.

